# Limb loss and specialized leg dynamics in tiny water-walking insects

**DOI:** 10.1101/2024.04.02.587762

**Authors:** Johnathan N. O’Neil, Kai Lauren Yung, Gaetano Difini, Pankaj Rohilla, M. Saad Bhamla

## Abstract

The air-water of the planet’s water bodies, such as ponds, lakes and streams, presents an uncertain ecological niche with predatory threats from above and below. As *Microvelia* move across the water surface in small ponds, they face potential injury from attacks by birds, fish, and underwater invertebrates. Thus, our study investigates the effects of losing individual or pairs of tarsi on the *Microvelia’s* ability to walk on water. Removal of both hind tarsi causes *Microvelia spp*. to rock their bodies (yaw) while running across the water surface at ±19°, compared to ±7° in non-ablated specimens. This increase in yaw, resulting from the removal of hind tarsi, indicates that *Microvelia* use their hind legs as ‘rudders’ to regulate yaw, originating from the contralateral middle legs’ strokes on the water’s surface through an alternating tripod gait. Ablation of the ipsilateral middle and hind tarsi disrupts directionality, making *Microvelia* turn in the direction of their intact limbs. This loss of directionality does not occur with the removal of contralateral middle and hind tarsi. However, *Microvelia* lose their ability to use the alternating tripod gait to walk for water walking on the day of contralateral ablation. Remarkably, by the next day *Microvelia* adapt and regain the ability to walk on water using the alternating tripod gait. Our findings elucidate the specialized leg dynamics within the alternating tripod gait of *Microvelia spp*., and their adaptability to tarsal loss. This research could guide the development and design strategies of small, adaptive, and resilient micro-robots that can adapt to controller malfunction or actuator damage for walking on water and terrestrial surfaces.

## Introduction

For tiny water-walking insects, venturing across the water’s surface involves more than balancing on surface tension. These tiny organisms encounter competition and predation from above, below, and on the water itself. Locomotion – an organism’s method of moving through its environment – serves as a significant evolutionary pressure shaping morphological traits [1]. In aquatic environments, the manner in which an insect moves across water often determines its vulnerability to predators, making the ability to quickly adapt to changing conditions essential for epineuston organisms living on the water surface. *Microvelia*, a water-walking insect that, unlike other water striders [2], possesses the ability to move on land [3], allowing it to navigate obstacles on the water’s surface, such as floating plants like duckweed, or even to flee to land to escape aquatic predators. Beyond maneuvering on the surface of water without sinking [4], water striders must contend with multiple predators in and out of the water [5], compete for resources and mates [6, 7, 8], and navigate the aftermath of conflicts that result in bodily damage. Should a *Microvelia* escape with its life but lose a limb, it faces the challenge of continuing to move on water.

Key evolutionary drivers for the ability to walk on water include predator avoidance, as seen in the basilisk lizard, mate displays or “rushing” in birds like Western and Clark’s grebes, and a combination of these factors for organisms that spend significant time at or near the water’s surface, such as fishing spiders (Dolomedes) or water-striders (Gerridae) [9, 10, 11]. For these smaller epineuston insects and spiders [3, 4], surface tension plays a crucial role in water locomotion. Their bodies, covered in hydrophobic wax and hairs, enables them to propel across the water without sinking [12, 13, 2, 14]. Although researchers have extensively studied the morphological adaptations that allow insects like *Microvelia* to walk on water [15, 16, 17, 18], and the unique use of the alternating tripod gait, similar to ants and cockroaches [19, 20, 21], the adaptations of *Microvelia* to limb loss remain unexplored. Adapting to limb loss is critical for insects like *Microvelia*, which cannot regenerate limbs after their final molt, necessitating their ability to compete on water despite limb absence. While insects commonly lose body parts [22, 23], and some may even shed limbs intentionally through autotomy [24, 25, 17], it becomes an additional challenge when every limb is the sink or swim determiner of difference in surface tension on the water.

In the wild, an injured *Microvelia* must generate sufficient force to walk on water without all its tarsi. The focus of this paper is on which leg’s removal affects direction and propulsion and how *Microvelia* adapts to limb loss. Previous studies have identified the middle legs as primary ‘propulsors’ due to their large stroke amplitudes compared to the front and hind legs [13], but the roles of other legs remain less understood. We explore the effects of tarsi loss on *Microvelia* locomotion by examining body velocity and directionality on water. Through high-speed imaging, pose estimation software (DeepLabCut) [26], and in situ ablation, we observe how *Microvelia*, despite these challenges, adapts and continues to navigate water surfaces.

## Materials and Methods

### Rearing and Experimental Setup

We collected *Microvelia* from ponds and creeks in Kennesaw, Georgia. The insects were housed in a 17.5 × 14.0 × 6.5 inch^3^ plastic container, filled with water maintained at a lab temperature of 20 °C, and supplemented with duckweed from their original habitats. We exposed the *Microvelia* to circadian lighting from 8 A.M. to 8 P.M. Additionally, from Monday through Friday, we fed the specimens daily with fruit flies procured from Carolina Biological. In total, we analyzed the locomotion of 20 specimens in response to the following ablations (N=3 specimens for each case): both front tarsi, middle tarsi, hind tarsi, both hind tarsi, contralateral middle and hind tarsi, and ipsilateral middle and hind tarsi. We observed only 1 specimen for both middle tarsi and single front tarsi ablations due to the lack of statistical significance in the single front tarsi ablation and the inability of *Microvelia* with both middle tarsi ablated to walk on water or survive beyond 48 hours post-ablation. Given these outcomes and the limited availability of specimens, we prioritized the preservation of specimens.

### *Microvelia* Tarsus Ablation

Before a particular *Microvelia* was ablated, it was anesthetized by placing it into a freezer for approximately 2 minutes. This led to the insect’s temporary incapacitation, which allowed for easier and more accurate ablation to be done. After being taken out of the freezer, we placed the *Microvelia* under a magnifying glass, and the according segment(s) (Fig1.c) of the leg(s) were removed with a Fine Science Tools dissecting knife. An example of a Microvelia with its middle tarsi ablated is shown in Fig1.a. After being cut, the specimen would reawaken and be placed into a small container of water from its natural habitat for recovery. After one hour of being in the container, the *Microvelia* was removed, its locomotion was recorded, and it was then placed back into containment. Additionally, after having 24 hours to recuperate from the initial ablation, the insect’s locomotion was once again recorded.

**Fig. 1.**
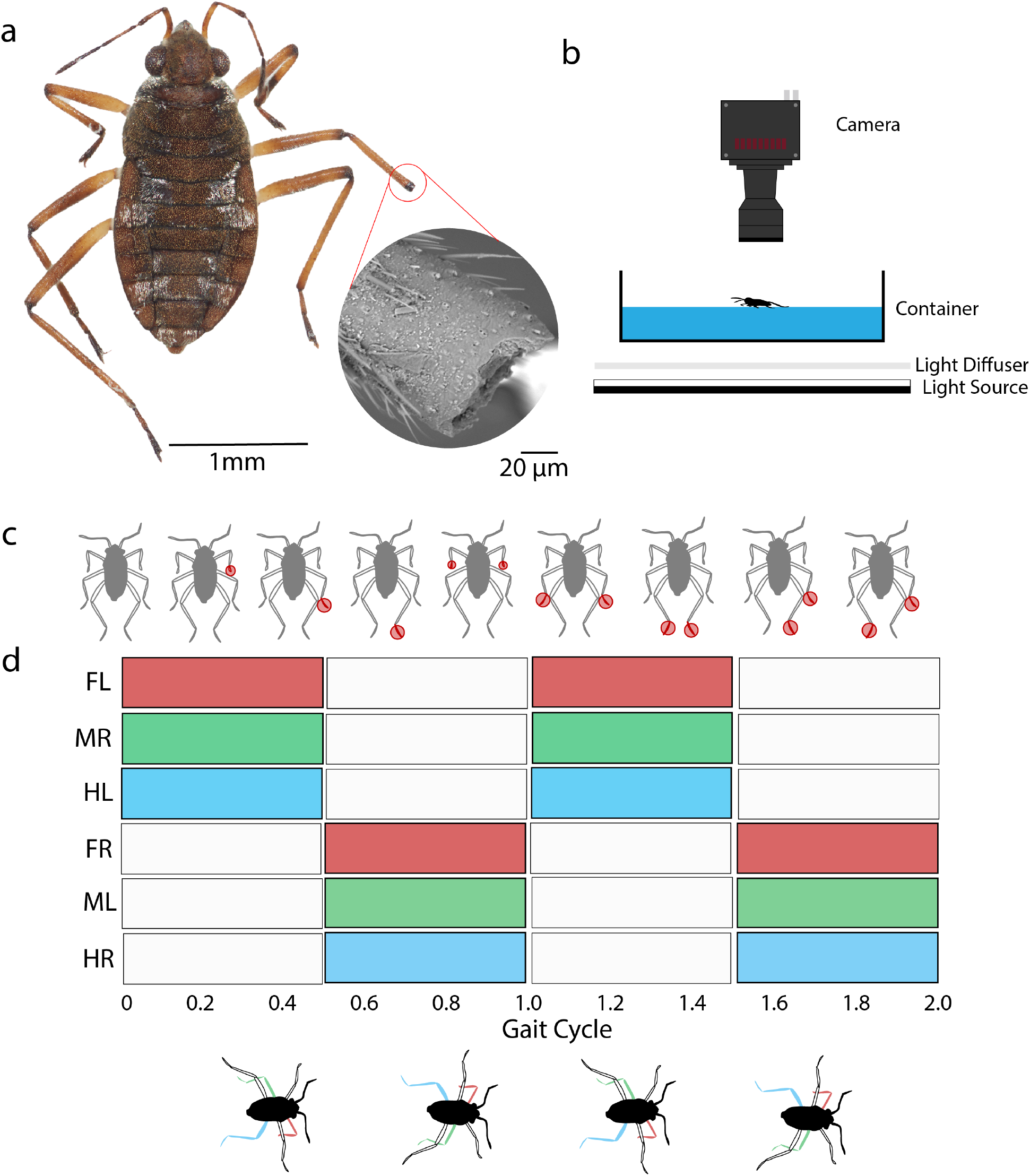
**(a)** High resolution z-stack image and Scanning Electron Microscopy image of a *Microvelia* with an ablated middle-right tarsi. **(b)** Schematic of experimental setup. A high speed camera is mounted above a container of water which rests on a diffuser. A light source is set at a short distance below the diffuser to provide even lighting when recording. *Microvelia* are recorded individually running on the water. **(c)** Illustration showing tested *Microvelia* ablations. Circled parts of legs indicate the different locations of ablations. Eight ablation conditions are investigated in this paper. **(d)** Gait cycle indicating the power stroke (colored rectangles) and recovery phase (blank rectangles) of the alternating tripod gait. Colored legs corresponding to the gait cycle showcase the alternating movement.

### Recording

To record the response of the *Microvelia* to their ablations, a Photron FASTCAM MINI AX 2000 was used with a frame rate of 1,000 - 2,000 frames per second at a resolution of 1024 × 1024 pixels. A Nikon 70-200mm f/2.8G ED VR II AF-S Nikkor Zoom lens was mounted onto the camera for enhanced documentation. The camera and was mounted vertically on a Thorlabs Optical Rail for a top view of *Microvelia* locomotion on water. The *Microvelia* were placed in a 10.0 × 10.0 × 1.5 cm^3^ petri dish (Thermo Fisher Scientific) that was filled halfway with water, and rested on top a white diffuser (Fig1.b). An LED light was also lit underneath the petri dishes for better lighting. The ablated insects were then prodded for movement, which was recorded and analyzed one video at a time.

### Tracking, Post Processing, and Analysis

After recording, DeepLabCut [26, 27] pose estimation machine learning software was utilized to track the head and abdominal tip of the *Microvelia* in each video. A custom Matlab script was used to calculate the displacement, velocity, and yaw angle from the DeepLabCut tracking data. For statistical analysis, we used one-way analysis of variance (ANOVA) test to find if the set of treatment effects yielded differences amongst the means of each group with post-hoc Tukey’s difference criterion to find which pairs of treatment effects were statistically different.

## Results

### Widened Yaw Angle

First, we track each specimen as it walks across the water surface (Fig2.b). In comparing non-ablated *Microvelia* to those *Microvelia* with both hind tarsi removed, we observe an increase in the rotation of the head along the movement direction. We then measure maximum body velocity of each specimen, based on the tarsi removed, and compare these velocities to that of the non-ablated specimens (Fig2.c). *Microvelia* with no tarsi removed achieve a maximum velocity (*v*_*max*_ = 14 cm/s, N=3 specimens, n=21 trials). Removing either both front or both hind tarsi yields no statistical difference in body velocity compared to non-ablated specimens (N=3, n=21, *p* > 0.05), which makes sense given that the middle legs are the main propulsors generating forward thrust. Consequently, removing both middle tarsi renders a *Microvelia* incapable of moving across the water (see Supplementary Movie S1), reducing its velocity to 2 cm/s. Next, we calculate the yaw angle over time for each tested specimen (Fig2.a). We find that, post-ablation, *Microvelia* exhibits an increase in yaw in both directions as they move on the water surface. Analyzing the yaw angle versus time data, we identify the delta yaw angle (Δ*θ* = *θ*_*f*_ − *θ*_*i*_) for each cycle, where the absolute value of the yaw angle is shown in (Fig2.d). For both non-ablated specimens and those with with front tarsi ablated, the yaw angle reaches Δ*θ* = ±7° as they run across the water surface (Fig2.d). Specimens with both hind tarsi ablated exhibit a yaw angle more than double (Δ*θ* = ±19°) that of the non-ablated and front ablated specimens (*p* < 0.001). This result underscores the role of the hind tarsi as ‘rudders’ that serve to minimize side-to-side rocking during water walking (see Supplementary Movie S2). We did not measure the yaw angle for specimens with both middle tarsi ablated as they cannot walk across the water surface.

**Fig. 2.**
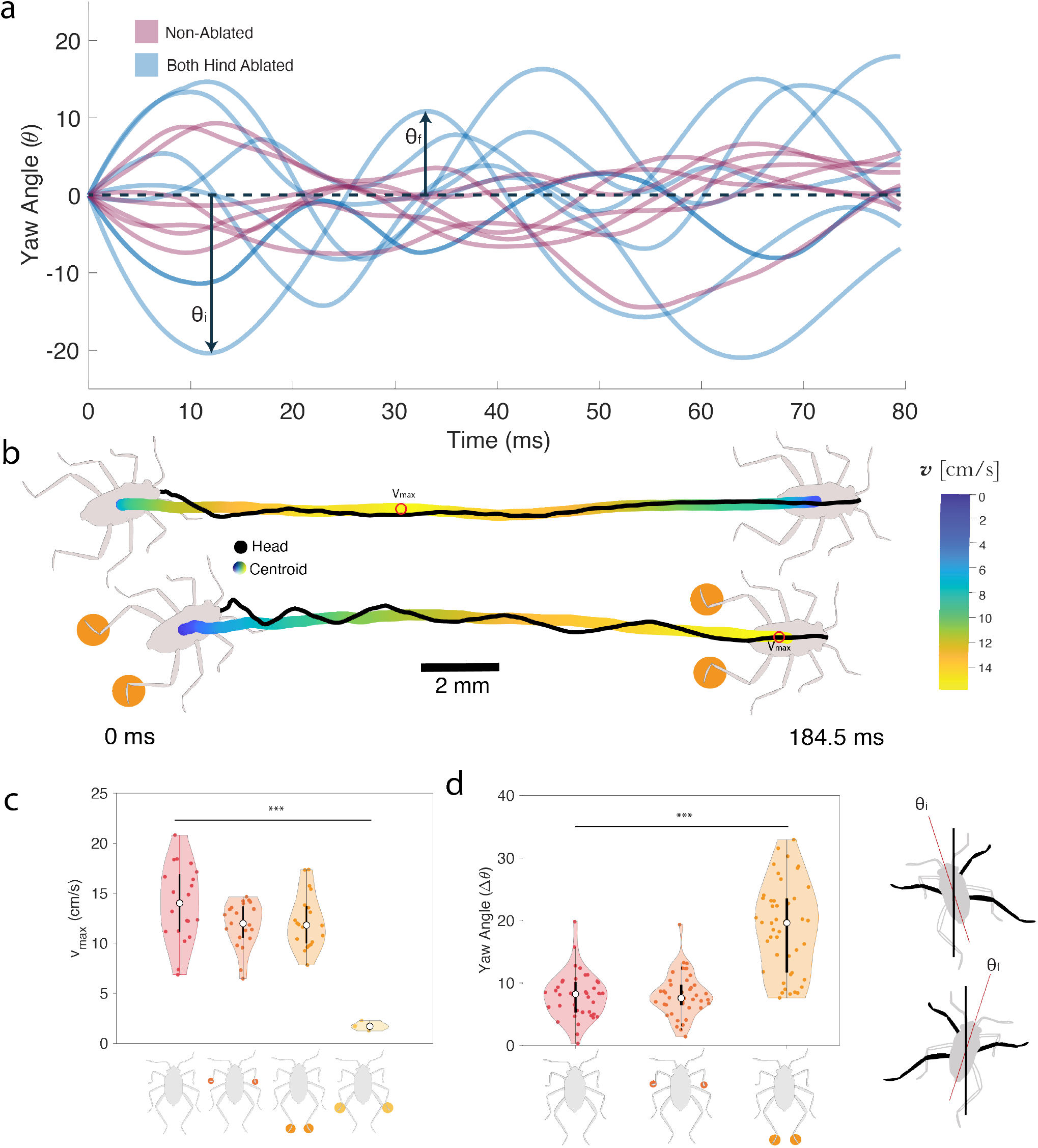
Yaw angle, trajectory, and maximum velocity for non-ablated and ablated *Microvelia*. **(a)** Yaw angle over time of a *Microvelia* before ablation (non-ablated) and after ablation (both hind ablated condition), with a visual difference in the size of yaw angles. **(b)** Trajectories of a non-ablated and both-hind ablated *Microvelia*. The increase in yaw of a both-hind ablated *Microvelia* is visibly greater. Red circles indicate the point of maximum velocity along the path. **(c)** Graph of maximum velocities (*v*_*max*_) of four *Microvelia* conditions. From left to right: control (non-ablated) *Microvelia* (N=3 specimen, n=21 trials), both front tarsi ablated (N=3, n=21), both hind tarsi ablated (N=3, n=21), and both middle ablated. *Microvelia* missing their middle tarsi lose the ability to propel themselves. Only one both middle ablated specimen was tested to preserve the population, since the specimen did not survive within 48 hours of ablation. The white circle represents the median. Other points represent experimental values from each trial. The box represents the second and third quartiles with the extended lines representing the first and fourth quartiles. **(d)** Graph of yaw angles (Δ*θ*) of three *Microvelia* conditions. From left to right: control (non-ablated), both front tarsi ablated, both hind tarsi ablated. When missing its hind tarsi, the *Microvelia*’s yaw angle increases. The yaw angle for both the left and right direction are plotted. p-value: ^*^ *p* < 0.05, ^**^ *p* < 0.01, ^***^ *p* < 0.001.

### Deviated Directionality

To assess the impact of tarsal loss on *Microvelia* directionality, we calculated the ratio of the final displacement (*D*_2_) to the total distance traveled (*D*_1_) for both non-ablated and ablated *Microvelia* (Fig3a). For a straight path, the ratio *D*_2_/*D*_1_ ∼ 1. For non-ablated specimens and most types of ablations, *Microvelia* typically travel in a straight line, with *D*_2_/*D*_1_ > 0.90. However, *Microvelia* missing ipsilateral middle and hind tarsi are the notable exception, exhibiting *D*_2_/*D*_1_ ≈ 0.86, indicating significant deviation from a straight line (*p* < 0.001). Fig3a illustrates this deviation with an example of an ipsilaterally ablated specimen traveling in a circular path and ending up facing the opposite direction from where it started (see Supplementary Movie S3). For the case of the contralaeral ablated *Microvelia* and the single tarsi ablated *Microvelia*, there is no significant deviation from the non-ablated specimen (*p* > 0.05) (see Figure 3.b).

**Fig. 3.**
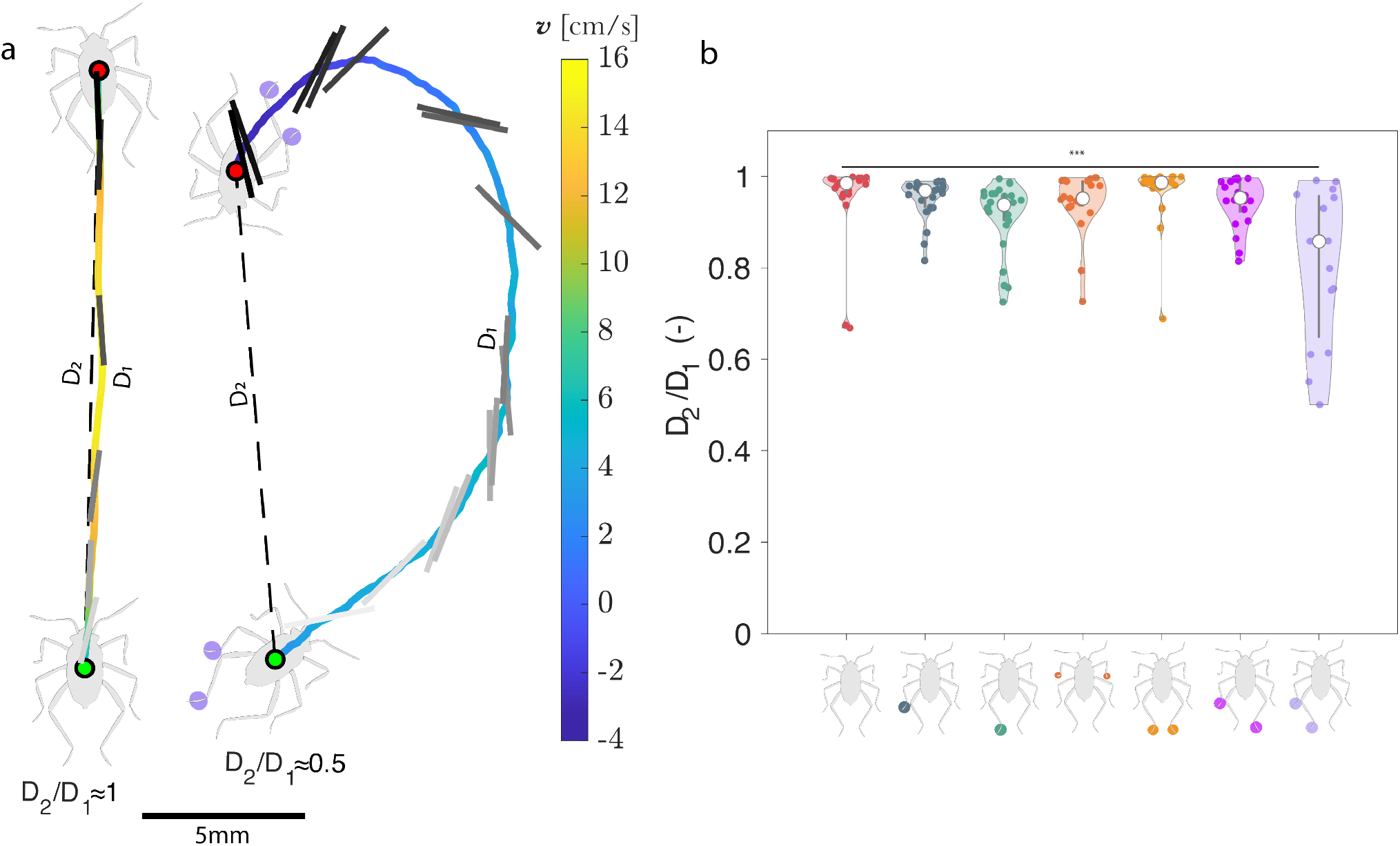
Displacement compared to actual distance travelled. **(a)** Displacement (*D*_2_) of *Microvelia* compared to distance travelled (*D*_1_), showcasing circular path for the ipsilateral ablated *Microvelia* vs. a non-ablated *Microvelia* which moves in a straight line. Green dot indicates the starting position, red dot indicates the ending. **(b)** Comparison of a set of *Microvelia* ablation conditions and their displacement-distance ratio travelled in each trial (N=3 for each ablation condition). White circle represents the median. Other points represent experimental values from each trial. Ipsilateral ablated *Microvelia* have a hindered directionality shown by its lower displacement-distance ratio (N=3,n=16). P-value: ^*^ *p* < 0.05, ^**^ *p* < 0.01, ^***^ *p* < 0.001.

### Adaptation to Tarsi Loss

*Microvelia* must adapt to the loss of key body locomotive parts (tarsi, leg) after ablation. We observed changes in the body velocities of *Microvelia* immediately following ablation (within an hour) compared to 24 hours later. Before ablation, *Microvelia* achieved a maximum body velocity (*v*_*max*_) of *∼*14 cm/s. Those missing their contralateral middle and hind tarsi initially struggled with a lower maximum velocity of 2 cm/s on the day of its ablation (N=3, n=15). However, by the next day their *v*_*max*_ increased to about 8 cm/s (N=3, n=21, *p* < 0.001, see Supplementary Movie S4). For *Microvelia* undergoing ipsilateral middle and hind ablation, despite also missing two tarsi, the difference in *v*_*max*_ between the day of ablation and the following day was not statistically significant (N=3, n=15, Fig4a, *p* > 0.05). The type of ablation also significantly affected their gait cycle. *Microvelia* with ipsilateral ablations continued to use the alternating tripod gait. In contrast, those with ablations on opposite sides displayed no discernible periodicity in their gait on the day of their ablation (Fig4b), yet managed to return to the alternating tripod gait within 24 hours (Fig4c).

**Fig. 4.**
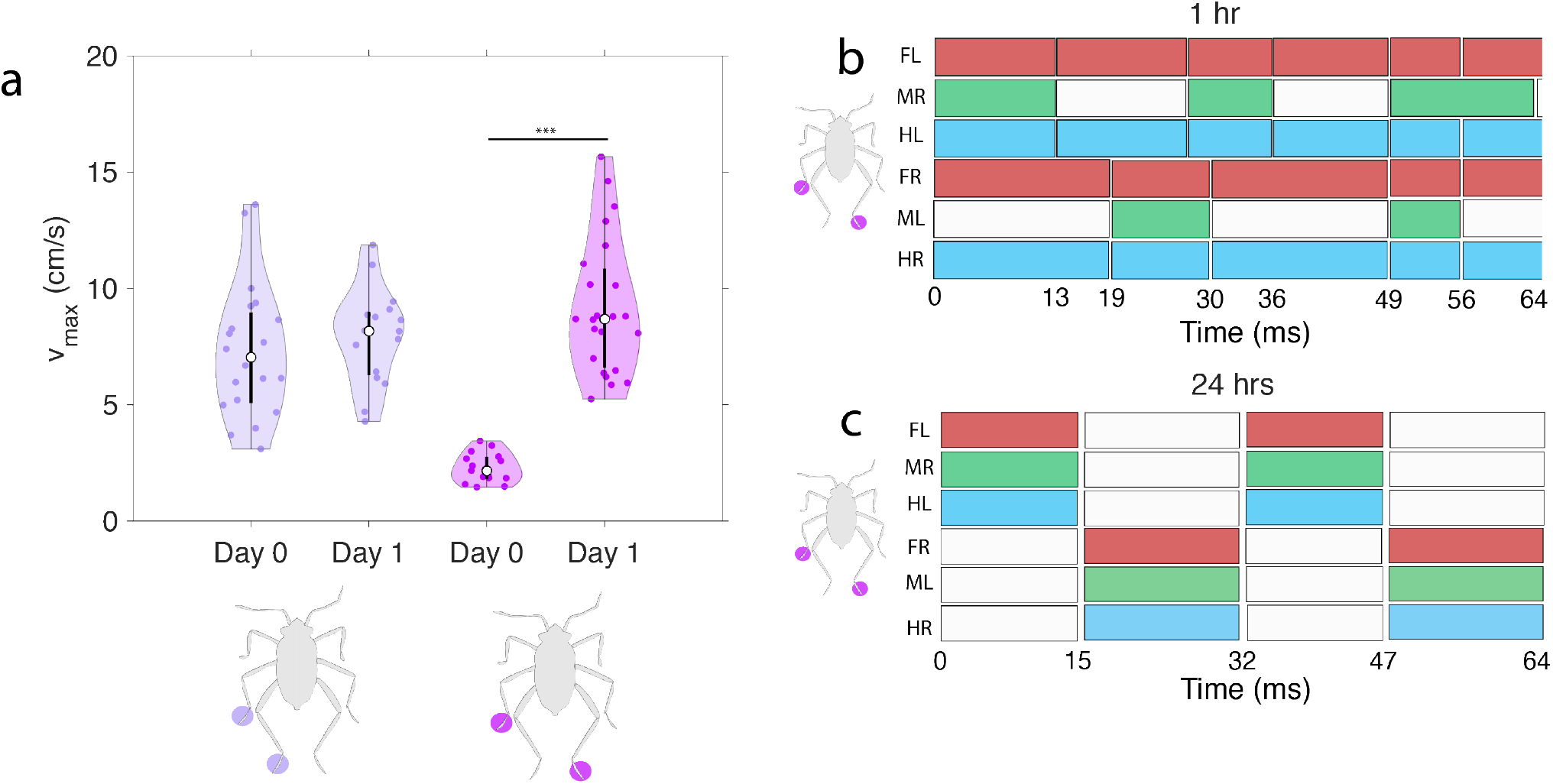
Velocity of *Microvelia* and how they adapt their gait. **(a)** Maximum velocity (*v*_*max*_) comparison of two ablated conditions, a contralateral middle and hind tarsi ablation and an ipsilateral middle and hind tarsi ablation (N=3 for both ablations), on the day they are ablated and one day after. For the contralateral ablation, Microvelia are unable to walk on water on the day of ablation, but can walk the next day. White circle represents the median. Lines represent the 1st and 4th quartile. Other points represent experimental values from each trial. The box represents the 2nd and 3rd quartile. **(b)** Gait plot of a *Microvelia* with a contralateral ablation within 1 hour of its ablation. **(c)** Gait plot of a *Microvelia* with an opposite side ablation > 24 hours after its ablation, which matches with the alternating tripod gait of non-ablated *Microvelia*. P-value: ^*^ *p* < 0.05, ^**^ *p* < 0.01, ^***^ *p* < 0.001.

## Discussion

For locomotion on the water surface, it is a well-documented strategy amongst water striders to rely on their middle legs as the primary propulsors of interfacial movement. Water striders such as *Gerridae* [13, 2, 4], *Rhagovelia* [28, 29], and *Velia* [13] use a rowing gait in which only the middle legs row against the water surface to propel themselves forward. The remaining legs are used only for support to float on the water’s surface. This research affirms that *Microvelia*, despite using an alternating tripod gait, prioritize their middle middle legs for propulsion, a finding consistent with previous studies [13]. The critical role of these legs becomes evident upon their ablation, which results in a significant decrease in velocity from 14 cm/s to 2 cm/s, underscoring their indispensability for water traversal.

Contrasting the alternating tripod gait of *Microvelia* with other hexapods that occasionally (or accidentally) enter aquatic environments reveals a unique adaptation in its locomotion strategy. For instance, *C. schmitzi* ants swimming in pitcher plant digestive fluids or ants that accidentally fall into water use both their front and middle legs for propulsion. Their front legs kinematically mimic terrestrial movement, their middle legs serve as rudders, and their hind legs act as roll stabilizers [19, 30]. Despite these ants employing an alternating tripod gait, their “swimming” episodes are brief and rare, lasting under 45 seconds in pitcher plant fluids or longer when they fall from a tree canopy into water. In contrast, *Microvelia* spends most of its time on water surfaces. The removal of *Microvelia*’s front legs does not impact velocity or directionality, highlighting their role in stability rather than in propulsion or direction. Ablation studies reveal that *Microvelia*’s hind legs function as rudders, facilitating directional movement and reducing yaw on water’s slippery surface.

The predator-prey dynamic underlines the importance of adaption for survival, not just in evading predators but in recovering from attacks. While many insects can regenerate limbs during larval stages after moulting [31, 32], many do not, especially after autotomy [24], muscle degeneration [33], or reaching final molting stages [34]. *Microvelia* undergoes five instars, after moulting ceases [35, 36], making any post-moult damage, such as tarsi or limb loss from a aerial bird or underwater fish, permanently affect their mobility and directionality.

*Microvelia* uses its middle legs as primary propulsors, causing a side-to-side rocking motion in the direction of the active leg (due to alternating leg strides). Without hind tarsi, this rocking motion intensifies, indicating their role as rudders (see Supplementary Movie S2). However, the removal of front tarsi does not alter the yaw angle. For a non-ablated specimen this is at a range of ±7°. Upon removal of both hind tarsi, the *Microvelia* rocks (yaws) at ±19°.

Our findings indicate that the extent and location of limb loss critically influence *Microvelia*’s ability to maintain direction while moving on water. Loss of both the middle tarsi is fatal as the organism cannot propel itself and eventually dies of fatigue. However, in most other cases of tarsi damage, *Microvelia* is still able to move on water after limb loss. Particularly, *Microvelia* with ipsilateral middle and hind tarsi removed show compromised straight-line movement, often veering off course (see Supplemental Movie S3). This impairment suggests challenges in predator evasion or prey capture due to reduced directional control.

Contrastingly, *Microvelia* contralateral middle and hind tarsi ablated initially lose the alternating tripod gait and show a significant drop in velocity, but within 24 hours, they regain the tripod gait (see Supplementary Movie S4) and approximate the speed of those with ipsilateral ablations, favoring straighter paths. These observations underscore the hind tarsi’s role in moderating yaw caused by contralateral middle leg movement, aiding in directional stability. This insight contrasts with the rowing gait, where any immobilization increases yaw [37], highlighting the alternating tripod gait’s biomechanical advantage in maintaining directionality despite limb loss. Thus, *Microvelia*, despite lacking the ability to regenerate limbs post-final molt, demonstrates remarkable resilience and adaptability in the face of physical impairments, adds another compelling narrative of survival and adaptation within the natural world.

Adaptable multi-surface gaits, such as the alternating tripod gait utilized by the *Microvelia*’s specialized leg dynamics, can be mimicked in future designs of small amphibious robots as much interest is gathering in robotics at the air-water interface and increasingly complex terrains [38, 39]. A robust robot will be able to traverse a variety of surfaces without having to enact more complex motion than an alternating tripod gait.

### Limitations and Future Outlook

Our study only focuses on the removal of tarsi as the removal of the femur or the entire leg would pose a greater impact on the overall balance, directionality, and velocity of the *Microvelia* which would conflate the roles that each leg has in locomotion. Our studies also use a relatively small sample size which may increase variation within our data and impact our results. Yet, our experimental setup was able to provide us consistent results. Future work can increase the sample size and also explore juvenile instars to further study the specialized dynamics of the *Mircovelia’s* legs and effect of organism size when walking on water. Future studies can also determine if other water walking insects that use the alternating tripod gait also have these specialized leg dynamics.

*Microvelia* has another means of propulsion on the water surface, Marangoni propulsion[13], in which it spits a fluid from its proboscis to lower the surface tension within a limited area. This reduction in surface tension allows the *Microvelia* to propel itself forward, and is used as an escape mechanism. Due to the reduced velocity cause by certain ablations, Marangoni propulsion can be a more prefered way to move in certain conditions such as predation. Future studies could study if the use of Marangoni propulsion is more likely when *Microvelia* is missing a tarsi or limb.

## Conclusion

Through ablation we investigate the specialized leg dynamics within the *Microvelia*’s alternating tripod gait. Through high-speed imaging and pose-estimation deep learning software, we measure the velocity, yaw angle, and directionality of the *Microvelia* with different missing tarsi. Our results show that *Microvelia* uses its hind legs as rudders to stabilize the direction of movement, while the middle legs are the main propulsers for locomotion on water. When the front legs were ablated, we observed no impact in overall body velocity or yaw angle, suggesting that the front legs help in stability when *Microvelia* walk on water.

When removing the contralateral middle and hind legs, the *Microvelia* was initially unable to traverse the water surface. Yet, the same specimen, adapted to their missing tarsi and performed the alternating tripod gait the next day. This contrasted with the removal of the ipsilateral middle and hind tarsi, where *Microvelia* were able to use the alternating tripod gait immediately after removal. However, *Microvelia* with the ipsilateral ablation had reduced directionality and sometimes traveled in curved paths rather than straight paths. These results suggest that the removal of middle and hind tarsi pose a threat to *Microvelia* in the wild as *Microvelia* would have higher difficulty avoiding predators or catching prey from their reduced body velocity and inhibited directionality. Ultimately, this study can influence the design of future robotics that may implement their own specialized leg dynamics for locomotion on the surface of water.

## Supplementary data

Supplementary data available at *ICB* online.

## Data Availability

The data underlying this article are available in the article and in its online supplementary material.

## Author contributions statement

J.O. designed the experiments. J.O, K.Y, and G.D. carried out the experiments and acquired the data. J.O. analyzed the data and interpreted the results. P.R. assisted with data analysis and manuscript review. M.S.B. reviewed the design and execution of experiments, data analysis, interpretations, and manuscript. All authors contributed to writing the manuscript.

## Acknowledgments

J.O. acknowledges funding support from the GT UCEM fellowshp program and the Herbert P. Haley fellowship program. M.S.B. acknowledges funding support from the NIH Grant R35GM142588 and NSF Grants CAREER 1941933 and 2310691. PR acknowledges funding support from them Eckert Postdoctoral Fellowship, Georgia Institute of Technology. We thank members of the Bhamla lab for useful discussions and feedback of this work.

