## Supplementary material for "Limb loss and specialized leg dynamics in tiny water-walking insects": Figures_SI _ICB_2024.docx

a

b
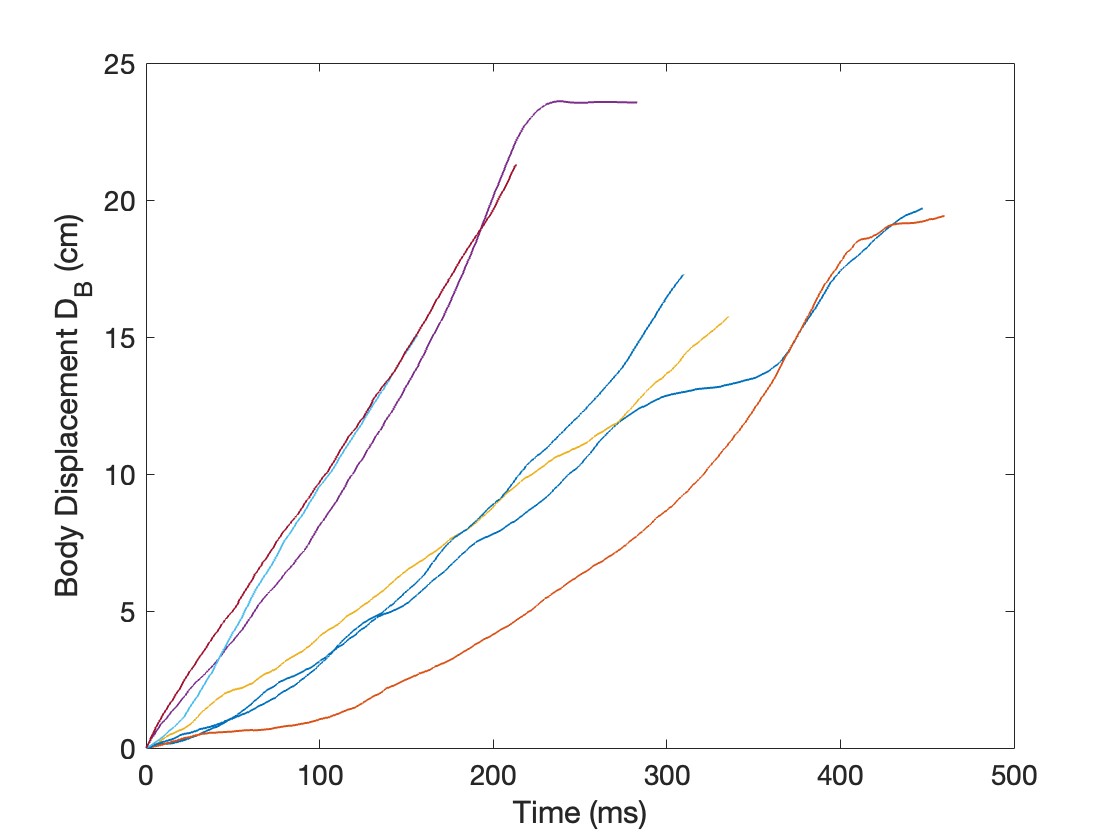

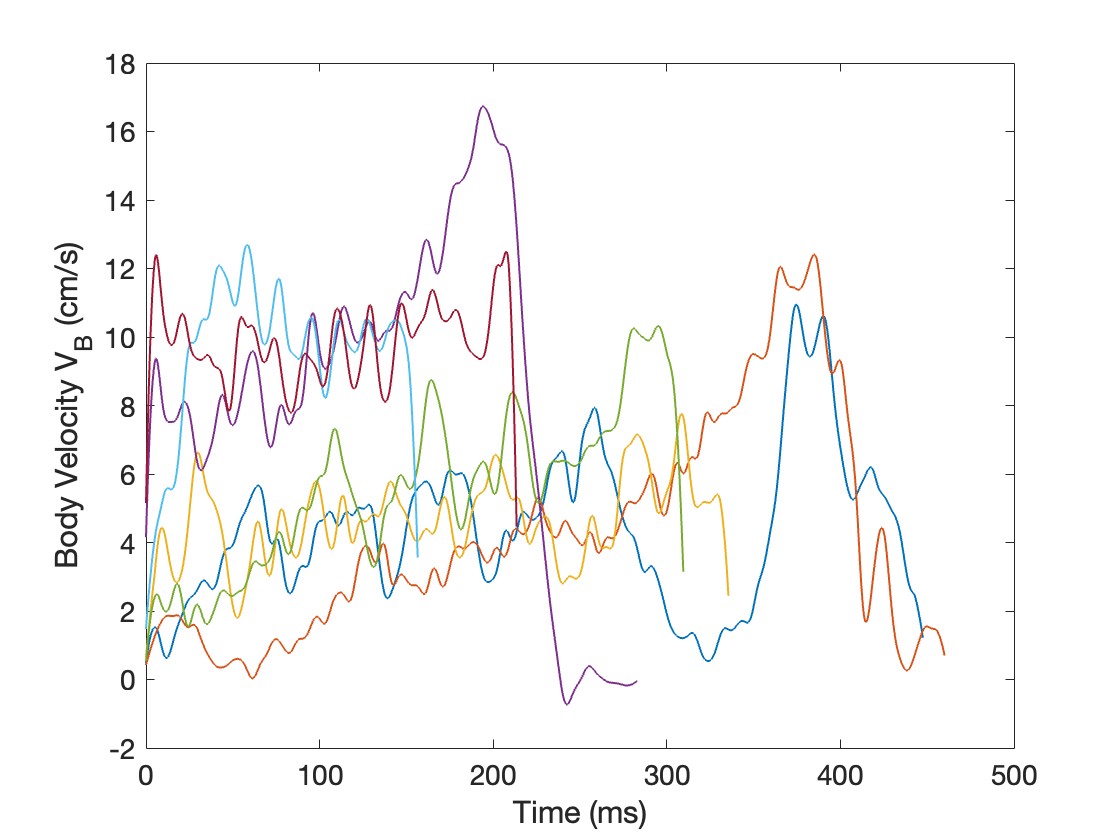

**Figure S1:** **Body Displacement and Body velocity over time for a Non-Ablated *Microvelia*** (a) Displacement versus time of a non-ablated Microvelia. (N=1 specimen, n=7 trials) (b) Velocity versus time of a non-ablated Microvelia. (N=1 specimen, n=7 trials)

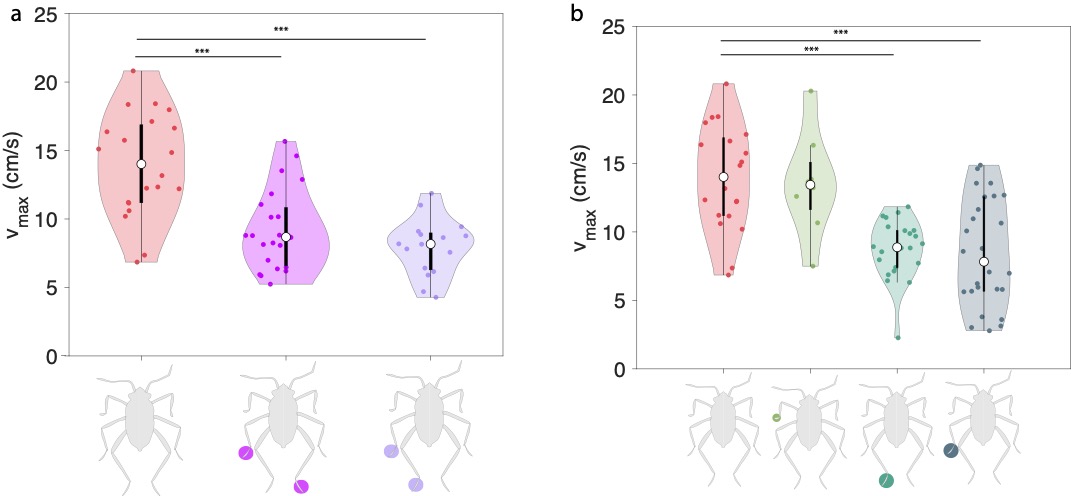

**Figure S2:** **Maximum body velocity for non-ablated and ablated Microvelia** (a) Maximum body velocities of non-ablated Microvelia (N=3 specimen, n=21 trials), contralateral middle and hind tarsi ablated Microvelia (N=3, n=21), and ipsilateral middle and hind tarsi ablated Microvelia (N=3, n=15). (b) Maximum body velocities of non-ablated Microvelia, front tarsi ablated Microvelia (N=1, n=5), hind tarsi ablated Microvelia (N=3, n=22), and middle tarsi ablated Microvelia (N=3, n=26). * p<0.05, ** p<0.01, *** p<0.001.

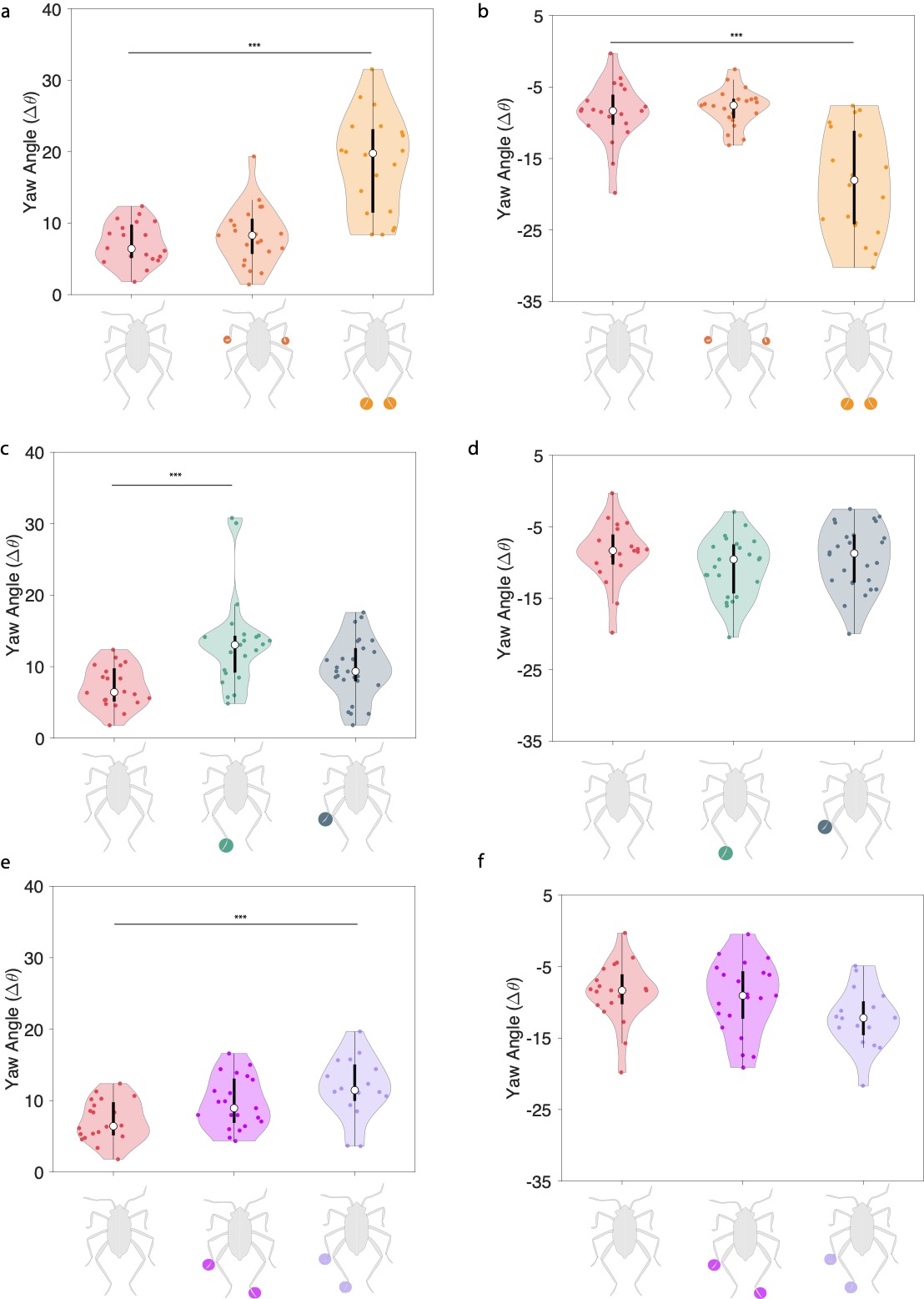

**Figure S3:**  **Yaw angle for non-ablated and ablated Microvelia** (a-b) Positive (right direction) and negative (left direction) yaw angle of non-ablated (N=3, n=21), both front ablated (N=3, n=21), and both hind ablated *Microvelia* (N=3,n=21). (c-d) Positive and negative yaw angle of non-ablated (N=3, n=21), middle ablated (N=3, n=26), and hind ablated *Microvelia* (N=3,n=22). (e-f) Positive and negative yaw angle of non-ablated (N=3, n=21), contralateral ablated (N=3, n=21), ipsilateral ablated *Microvelia* (N=3, n=16).
